# Dual Energy X-ray Absorptiometry (DEXA) as a Longitudinal Outcome Measure of Cancer-Related Muscle Wasting in Mice

**DOI:** 10.1101/2020.03.09.983403

**Authors:** Calvin L. Cole, Deja Robinson, Jian Ye, Bradley Mills, Scott A. Gerber, Christopher A. Beck, Edward M. Schwarz, David Linehan

## Abstract

**Introduction:** Pancreatic ductal adenocarcinoma (PDAC) is notorious for its associated skeletal muscle wasting (SMW) and mortality. Currently, the relationships between PDAC, SMW, and survival are poorly understood. Thus, there is a great need for a faithful small animal model with a quantitative longitudinal outcome measure that recapitulates clinical PDAC, to define SMW onset and assess progression. Therefore, we aimed to validate dual energy X-ray absorptiometry (DEXA) as a longitudinal outcome of lean body mass, and demonstrate its utility to quantify SMW in the KCKO murine model of PDAC.

**Methods:** In vivo body composition of: 1) untreated mice at 5, 8, 12, 18, and 22 weeks of age (n=4), and 2) a cohort of mice with (n=20) and without PDAC (n=10), was determined via DEXA, and lean mass of the lower hind limbs was predicted via a region of interest analysis by two independent observers. Total body weight was determined. Tibialis anterior (TA) muscles were weighed and processed for histomorphometry immediately post-mortem. Statistical differences between groups were assessed using t-tests and ANOVA. Linear regression models and correlation analysis were used to measure the association between TA and DEXA mass, and reproducibility of DEXA was quantified via the intraclass correlation coefficient (ICC).

**Results:** Lean mass in growing untreated mice determined by DEXA correlated with TA mass (r^2^ = 0.94; p <0.0001) and body weight (r^2^ = 0.89; p <0.0001). DEXA measurements were highly reproducible between observers (ICC = 0.95; 95% CI: 0.89-0.98). DEXA and TA mass also correlated in the PDAC cohort (r^2^ = 0.76; p <0.0001). Significant SMW in tumor-bearing mice was detected within 38 days of implantation by DEXA, TA mass, and histomorphometry.

**Conclusions:** DEXA is a longitudinal outcome measure of lower limb lean mass in mice. The KCKO syngeneic model is a *bona fide* model of PDAC-associated SMW that can be quantified with longitudinal DEXA.

## Introduction

Pancreatic ductal adenocarcinoma (PDAC) is the most common malignancy of the pancreas, and is the fourth leading cause of cancer-related deaths, with its incidence expected to increase over the coming decade (1, 2). Despite advances in the treatment of PDAC, the 5-year survival rate remains below 10% (3). Additionally, treatment intolerance and/or discontinuation of treatment continue to present challenges for PDAC patients and caregivers. Most notable among the detractors of quality of life for PDAC patients is sarcopenia, also known as skeletal muscle wasting (SMW), which is a growing burden among cancer survivors (4, 5). As such, SMW is prognostic of treatment failure, radiotherapy toxicity, and a shorter time to tumor progression related to survival (6–9). SMW can be defined as any loss of muscle tissue, function, and/or strength due to aging, chronic diseases, low protein-energy intake and physical inactivity (10). Importantly, a large percentage of patients with PDAC experience cancer-related SMW (10), and these patients have reduced physical function, increased postoperative morbidity, reduced response to chemotherapy, and shorter life expectancy (11). Furthermore, SMW has been identified as a prognostic factor in pancreatic cancer (12) and is an independent predictor of infectious disease and postoperative mortality in resected patients (13),(14). Thus, reductions in the incidence of SMW in patients with pancreatic cancer may reduce disease and treatment-related complications, which adversely affect the dose and length of treatment.

Currently, the time of onset of SMW following a cancer diagnosis is poorly understood. However, clinical studies suggest that once cancer-related SMW is initiated, it is irreversible (15). Therefore, a priority in the treatment of PDAC-related muscle wasting must be determining when it is initiated and preventing its establishment. Clinically, this is relevant because identifying the onset of cancer-related SMW in an animal model may lead to the discovery of paracrine factors that are emitted in the early stages of disease that perpetuate SMW. The anatomical distance between tumor cells and sites of SMW posit that inflammatory cytokines may transmit systemic signals that potentiate muscle wasting through the alteration of myofibrillar intracellular pathways regulated by both hormones and cytokines that slow protein synthesis and accelerate catabolism (16). Unfortunately, further elucidations of PDAC-related SMW and identification of treatable targets have been challenging due to the absence of small animal models with longitudinal outcomes.

Early recognition of SMW may also aid clinicians in devising an appropriate dosing algorithm to reduce treatment toxicity, while improving treatment tolerance and related outcomes of the cancer diagnosis. In the context of non-metastatic disease, the only possible cure for PDAC is surgical resection (17). However, less than 20% of PDAC patients meet the criteria for resection due to the locally advanced or metastatic nature of their diagnosis (9). Recent research suggests that neoadjuvant treatment (NAT) may help to improve the resectability rates among patients with PDAC, but may have an adverse effect on body composition that worsen post-surgical outcomes or reduce resection opportunities (9). Among many other reasons, the adverse effect of NAT may be the result of an incorrect dosing regimen that is based on body weight or body mass index (BMI). Indeed, studies have shown the measurement of lean mass to be a superior indicator of treatment toxicity and dosing response (18) in patients who experience cancer-related SMW, when compared to body weight and BMI. Therefore, it may be beneficial to monitor a patient’s body composition before, during, and after neoadjuvant treatment to determine the early need for additional intervention.

Dual-energy X-ray absorptiometry (DEXA) has emerged as a viable, non-invasive method of serial in vivo body composition analysis in small animals, due to its feasibility, accuracy, and reproducibility(19). Additionally, DEXA is widely used for body composition measurements in humans for both clinical and research purposes. Thus, DEXA outcomes in an animal model of disease may recapitulate clinical outcomes. Furthermore, use of a non-invasive form of lean mass (LM) measurement is vital during PDAC-related SMW to better understand the onset of this phenomenon. In a homeostatic myocellular environment, pathways that regulate protein synthesis and breakdown function to prevent unnecessary protein cycling. However, in an environment of SMW, a dysregulation of anabolic and catabolic systems exists that results in a net loss of protein. A determination of the timing of this dysfunction may play a key role in understanding the mechanism(s) that lead to PDAC-related SMW. Therefore, we aimed to validate dual energy X-ray absorptiometry (DEXA) as a longitudinal outcome of lean body mass, and demonstrate its utility to quantify SMW in the KCKO murine model of PDAC (20).

## Materials and Methods

### Experimental Model

#### Aged Mice

The University of Rochester Medical Center University Committee on Animal Resources (UCAR) has approved of all animal work conducted herein. Female C57BL/6 mice were purchased from the Jackson Laboratory (stock number 000664) at 4, 7, 11, 17, and 21 weeks of age. Mice were aged in a pathogen-free facility for one week, under an IACUC approved protocol. Anesthesia and Euthanasia were performed in accordance with our approved protocol. Briefly, on the day of sacrifice, mice were anesthetized using 10μl of ketamine per gram of bodyweight. After sedation, mice underwent a full body DEXA scan. Mice were sacrificed immediately after the DEXA scan and their lower hind limb muscles were harvested for analysis.

#### Murine orthotopic model of pancreatic cancer

We utilized the murine syngeneic-orthotopic model of PDAC as previously described (20). As there is no known sexual dimorphism in this model, only female C57BL/6J mice were used. They were obtained at 6-8 weeks of age from the Jackson Laboratory (stock number 000664) and maintained in a pathogen-free facility under an IACUC approved protocol. As the development of this model has been described elsewhere (20), we will only briefly describe the model. After a 1-week acclimation period, mice were randomized to one of two groups: PDAC (n=5) or no tumor control (NTC) (n=5). Mice in the PDAC group were anesthetized and injected in the tail of the pancreas with 2×10^5^ KCKO-luc cells suspended in a 1:1 PBS to Matrigel (Corning) mixture. NTC mice received no surgery and were sacrificed at a ratio of 1:1 with mice of the PDAC group. Mice were maintained in standard isolation cages with a 12hr light: dark cycle, and given ad libitum access to water and standard chow. Longitudinal DEXA scanning was performed on days 14, 35, 42, 49, and 56. The total length of this experiment was 56 days. Tumor-bearing mice were sacrificed when they developed end-stage disease defined by exhibiting three or more characteristics defined by the Institutional Animal Care and Use Committee (IACUC) as “failure to thrive”. Characteristics of “failure to thrive” included, but were not limited to self-isolation, hunched over appearance, lack of or reduced cage activity, lack of or no resistance to scruffing, mangled hair appearance after scruffing, failure to eat or drink, and/or visual signs of breathing difficulty. To determine if these characteristics were being exhibited, animals were checked twice daily, 30 days after tumor inoculation. This is the time point in which untreated animals reach an advanced stage of disease (20). Mice in the non-tumor-bearing group were sacrificed concordantly to allow for intergroup comparisons. Following euthanasia, skeletal muscles (quadriceps, Extensor Digitorum Longus (EDL), Soleus (SOL), and Tibialis Anterior (TA)) and PDAC tumors were harvested for histology, and cardiac puncture was performed to obtain serum for Luminex assay.

#### Dual-energy X-Ray Absorptiometry (DEXA)

Body composition was assessed in all mice using a DEXA scanner (PIXImus2; Lunar, Madison, WI). Each mouse was anesthetized for the duration of the procedure (5 min) with an i.p. injection of 100 mg/kg ketamine. Each mouse was placed on the scanner bed in the prone position, with the limbs and tail stretched away from the body. The PIXImus employs a cone beam X-ray source generating energies at 35 and 80 keV with a current of 0.5 mA for both energy levels. The detector is flat (100 × 80 mm) and comprised of individual pixels of 0.18 × 0.18 mm. Based on the attenuation of two energy levels, the system provides quantitative data on the fat tissue content, the lean tissue content, and the total tissue mass within the region of interest (ROI). One scan per mouse was performed and analyzed with PIXImus software (2.10; GE/Lunar). The head was excluded from calculation using a manual ROI. The PIXImus was calibrated with an aluminum/lucite phantom (corresponding to bone mineral density = 0.0592 g/cm2 and 12.5% fat) on each day of testing according to the manufacturer’s instructions. Lean mass was calculated using the lower hindlimbs as an ROI to exclude the measurement of the tumor burden. Lean mass was calculated as an index of total mass minus fat mass using the following equation: total mass – ((% fat x total mass)/100). For each mouse, the lean mass was calculated for the lower right and left limb independently, and the average of both measurements was used as the final lean mass for the animal.

#### Antibodies

The following antibodies were used: laminin (rat, 1:1500, Sigma-Aldrich, L0663) and DAPI (1:3000).

#### Muscle Histology

TA, SOL, and EDL muscles were harvested for histology. TA muscles were cut from the most distal and proximal TA tendon attachment, cleaned of extraneous tissue, blotted, and weighed. After which, TA, muscles were stored in a 30% mixture of sucrose in PBS for 24 hours, at which time, muscles were embedded in OCT (Tissue Tek), flash-frozen using dry ice and 2-methylbutane (Sigma-Aldrich), and processed for fresh-frozen histology as previously described (21). 10μm sections were cut and stained with H&E and representative micrographs were obtained for descriptive analyses.

Immunohistochemistry for laminin (extracellular matrix) and nuclei determination were also performed as previously described (21, 22), in which TA muscles were cryosectioned at 10 μm, to obtain transverse sections. Muscle sections were fixed in 4% paraformaldehyde (PFA) for 10 min. Sections were permeabilized with PBS-T (0.2% Triton X-100 in PBS) for 10 min, blocked in 10% normal goat serum (NGS, Jackson ImmunoResearch) for 30 min at room temperature. If a mouse primary antibody was used, sections were blocked in 3% AffiniPure Fab fragment goat anti-mouse IgG (H+L) (Jackson ImmunoResearch) with 2% NGS in PBS at room temperature for 1 h. Primary antibody incubation was performed in 2% NGS/PBS at 4°C overnight. Secondary antibody incubation was carried out in 2% NGS/PBS at room temperature for 1 h. After washing in PBS, sections were counter-stained with DAPI to label myonuclei. All slides were mounted with Fluoromount-G (SouthernBiotech). Fluorescent microscopy was performed using a Zeiss Imager: M1m microscope with AxioVision SE64 software, and representative images were used to quantify muscle cell area.

#### Fixed Single Fiber Staining

Single myofiber size and myonuclear analysis was performed, as previously described (21). For single myofiber size and myonuclear analysis, whole limbs (n=10) were fixed in 4% PFA for 48 h prior to EDL and SOL muscle dissection. Fixed muscles were incubated in 40% NaOH for 2 h to induce dissociation, and single myofibers were gently titrated and washed in PBS prior to staining with DAPI. For quantification, the cross-sectional area (CSA) of 100 fibers per mouse was determined manually from digitally-photographed DAPI stained fibers at 100X, and averaged as the CSA for the mouse.

#### Statistical Analysis

Myofiber CSA was determined using ImageJ software. The diameter of the fiber was measured at three points along the fiber to get an average CSA. TA cell area was calculated by drawing an ROI around the extracellular space of 200 individual cells and taking the average of the sum of those cells. Results are presented as mean±sd. Statistical significance was determined using t-tests for single comparisons and one-way and two-way ANOVA for multiple comparisons. Pearson correlation coefficient was calculated to measure the association between TA weight vs. DEXA lean mass and body weight vs. DEXA lean mass. Linear regression models were used to evaluate DEXA as a predictor of TA weight and body weight with and without adjustment for the effects of time (treated as a categorical covariate to avoid assuming linearity over time), with predictability measured by the model’s square root of mean squared error (rMSE). Reproducibility of DEXA was assessed via the intraclass correlation coefficient (ICC) along with its 95% CI. Analyses were performed using GraphPad Prism software (GraphPad Software, San Diego, CA, USA) version 7.2 and 8.0, R version 3.5.1, and SAS version 9.4. P<0.05 was considered significant (*P<0.05, **P<0.01, ***P<0.001, ****P<0.0001).

## Results

### In vivo DEXA measures are reliable and reproducible in predicting the body mass of growing mice

To assess the utility of longitudinal DEXA to assess in vivo murine body mass, we completed DEXA scans on 5, 8, 12, 18, and 22 week old mice (Figure 1). DEXA revealed a significant increase in lower limb lean mass of growing mice from week 5 to week 18 (Figure 2A). Lower limb mass was measured by two independent observers and the reproducibility of DEXA proved to be excellent. The intra-class correlation coefficient (ICC) analysis completed on these measures shows strong agreement between the two separate observations (Figure 2B). TA and body weight analysis confirmed the DEXA results, as both averages increased with age in this cohort of growing mice (Figure 2C,E). In addition, regression modeling revealed substantial variation due to time, such that adding time as a covariate improved the predictability of lean mass by DEXA. Furthermore, this analysis determined that DEXA has an rMSE of 1.89 mg when predicting TA weights. Likewise, a Pearson correlation analysis demonstrated a strong relationship between DEXA and TA and body weight (Figure 2D,F).

**Figure 1:**
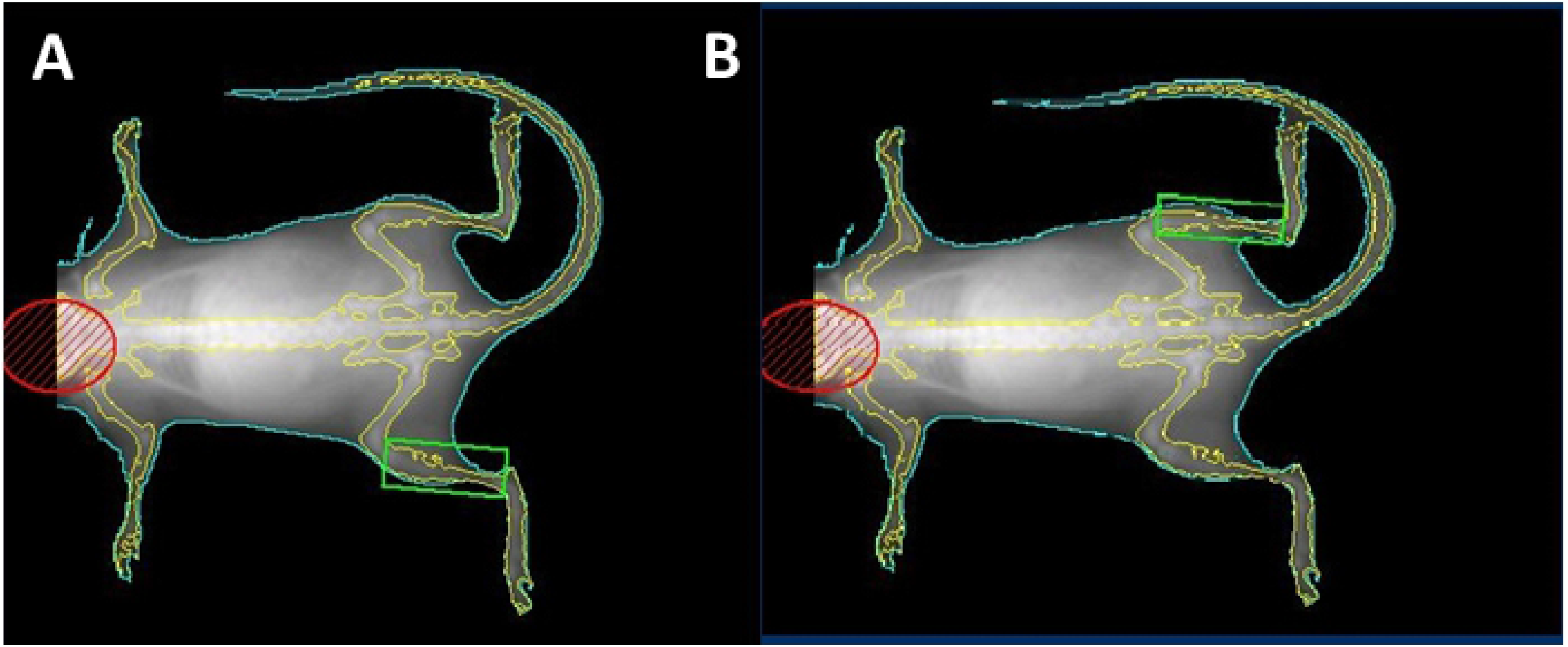
In vivo visualization and quantification of lean mass in a lower limb region of interest (ROI). A representative dual energy x-ray absorptiometry (DEXA) scan image of a mouse is shown to illustrate segmentation of the head (red oval), body (green outline), and the ROI for lean mass analysis (green box) of the (A) right and (B) left lower hind limbs as described in Material and Methods.

**Figure 2.**
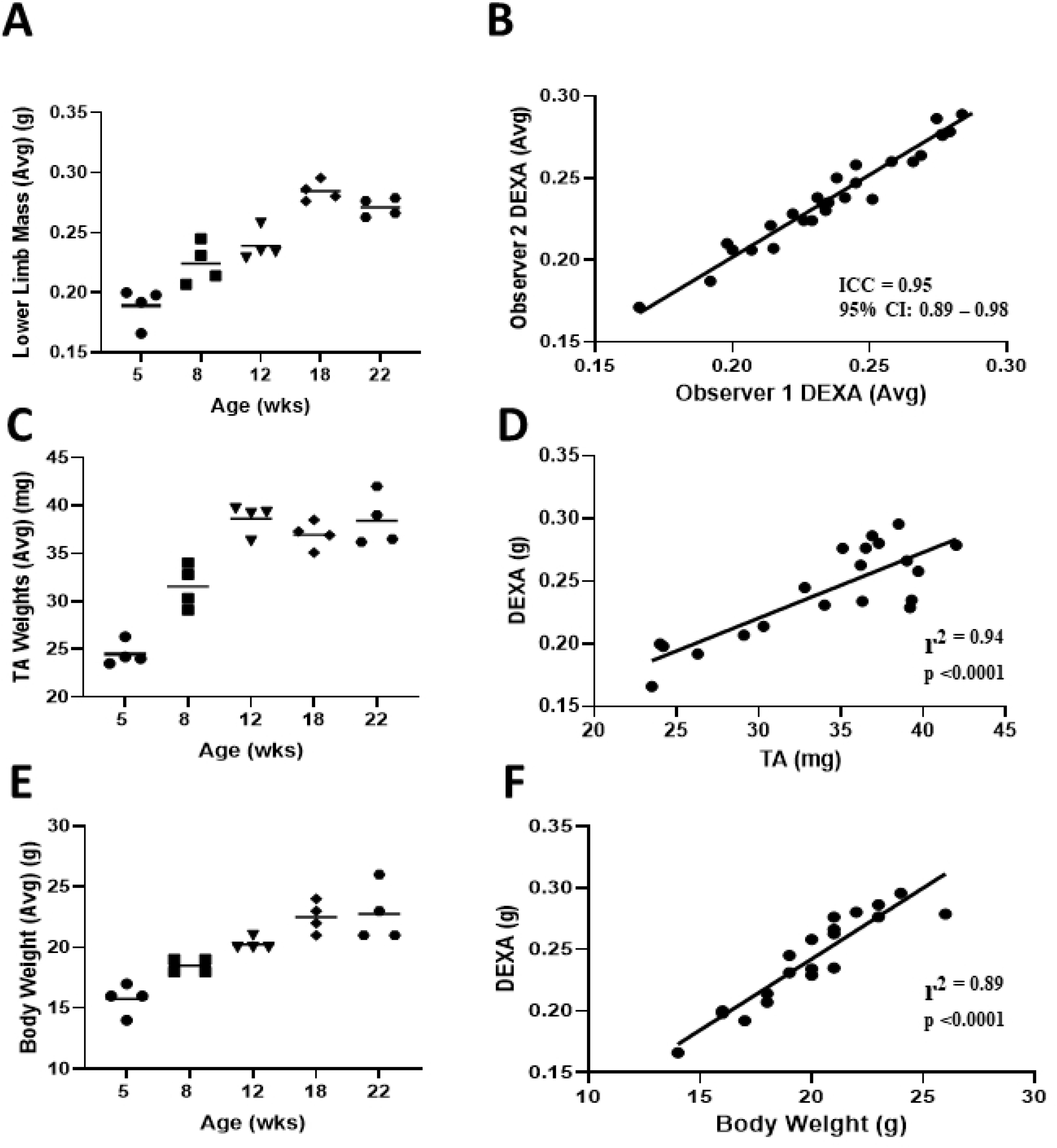
Validation of DEXA as an in vivo measure of lean mass in lower limbs of growing mice. Dual energy X-ray absorptiometry (DEXA) was performed on female mice at 5, 8, 12, 18 and 22 weeks of age (n=4 per group), postmortem, mice Tibialis Anterior (TA) muscles were dissected and weighed immediately. (A) The cross-sectional lower limb mass for each mouse was determined by DEXA at each time point by two independent observers as described in Figure 1, and the average mass of the two readings is presented with the mean for each group. (B) The intraclass correlation coefficient (ICC) was determined to assess the between reader variability of the DEXA measurements. (C) average TA weights of each group is presented as the mean ±SD. (D) A Pearson correlation analysis was performed to determine the relationship between DEXA lean mass vs. TA weight. (E) Average body weight of each group is presented as the mean ±SD. (F) A Pearson correlation analysis was performed to determine the relationship between DEXA lean mass vs. body weight.

### DEXA predicts skeletal muscle wasting in a murine model of PDAC

To assess the onset and progression of SMW in PDAC-bearing mice, we performed longitudinal, in vivo DEXA analyses normalized to total body weight (TBW) (weight including the tumor), and compared the findings to those in no tumor controls (NTC). DEXA revealed a decrease in the lower limb mass of tumor-bearing mice beginning at day 38, resulting in a significant decrease vs. NTC mice at day 56 (Figure 3B), a decrease that could not be detected by the measurement of TBW (Figure 3A).

**Figure 3.**
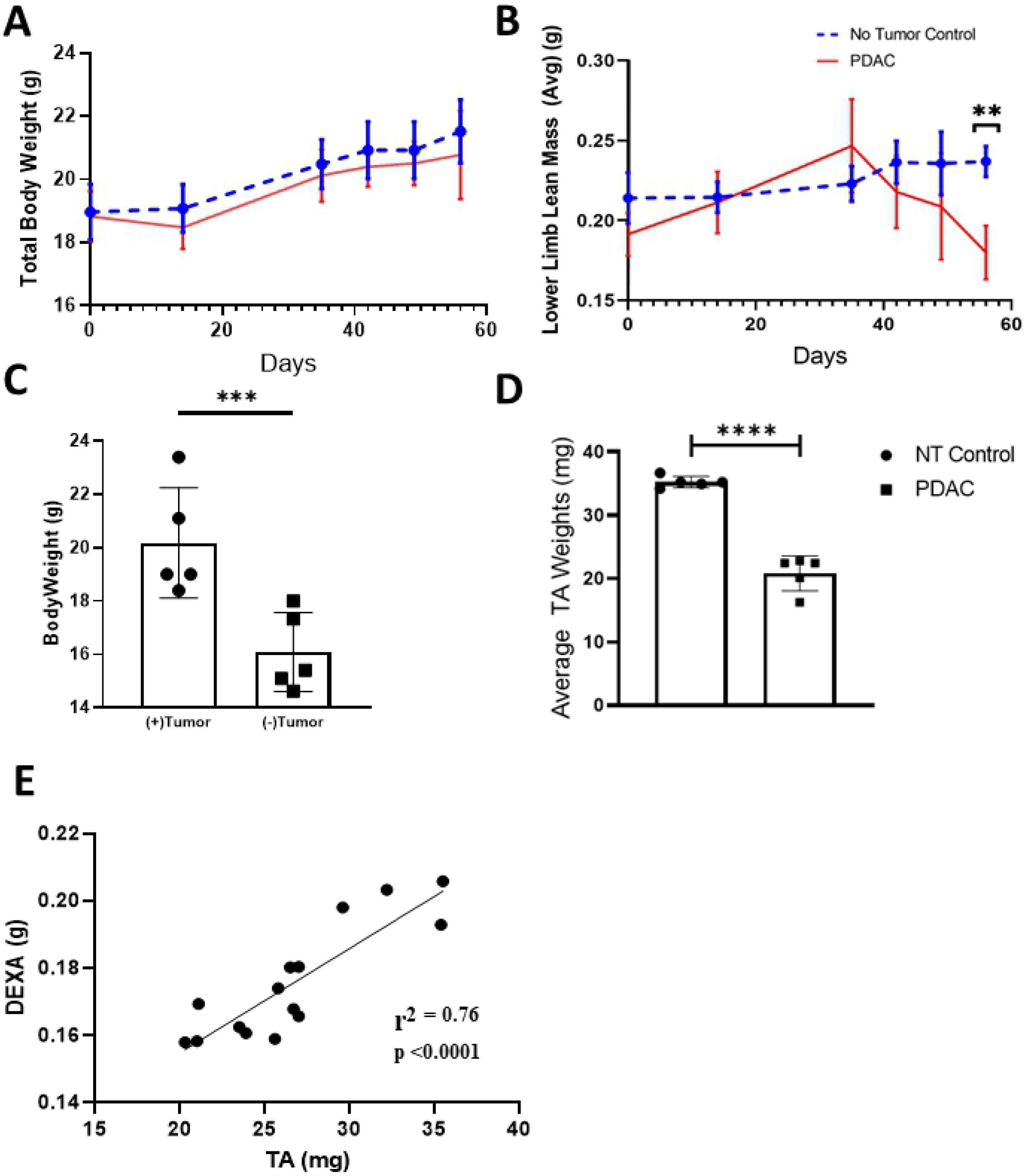
Longitudinal quantification of lean mass via DEXA demonstrates commencement of PDAC-related muscle wasting within 38 days of tumor implantation, which cannot be detected by assessment of total body weight. Mice were randomized to two groups prior to orthotopic tumor injections: 1) NTC; (n=5), and 2) PDAC tumor bearing; (n=5). Longitudinal DEXA and total body weight (TBW) measurements were performed as described in Materials and Methods. (A) Longitudinal TBW is presented for the NTC and PDAC mice that were sacrificed on day 56. (B) Longitudinal lower limb mass determined by DEXA is presented (**p<0.01). (C) The total body weight (TBW) (g) and (D) TA weights (mg) of the PDAC bearing mice was determined on the day of sacrifice, and the data are presented as TBW (+ tumor) before primary tumor harvest and net weight (− tumor) after primary tumor harvest for each mouse with the mean +/− SD (***p=0.0005) (****p<0.0001. (E) A Pearson correlation analysis was performed on a separate and larger cohort of PDAC tumor bearing mice (n=15) to determine the relationship between DEXA lean mass and TA weight.

Postmortem examination of each animal determined that TBW is an unreliable measure of the onset of SMW because the growing tumor mass compensates for the loss of viable tissue. Thus, the TBW of the tumor-bearing mice was significantly greater than their net weight (NW) (weight – tumor weight) (Figure 3C), and this increase was commensurate with primary tumor weight, such that there were no differences in TBW between the PDAC and NTC groups. Smaller TA muscles, as determined by weight, in the tumor-bearing mice when compared to the NTC group is further evidence of PDAC-associated SMW (Figure 3D). A Pearson correlation analysis, performed on a larger cohort of PDAC mice (n=15), confirmed a strong relationship between DEXA lean mass and TA weight (Figure 3E) in this cohort of mice as well.

### Immunohistochemistry confirms PDAC-related SMW

To confirm SMW is this model of PDAC, we performed histomorphometry on fast twitch (EDL) and slow twich (SOL) skeletal muscle fibers from the PDAC-bearing and NTC mice (Figure 4A,B). We found a significant decrease in the CSA of both muscle fiber types in the tumor-bearing mice, and that the 38% and 33% decrease is similar to reported decreases in EDL and SOL muscles respectively, in mice with age and disease-related sarcopenia (23). Histomorphemetry of TA sections (Figure 4E-H) show an atrophied appearance, while quantification of the CSA of these muscle fibers (Figure 4I) substantiate SMW. Collectively, these results formally establish SMW in this PDAC model, which can be longitudinally assessed via DEXA scanning.

**Figure 4:**
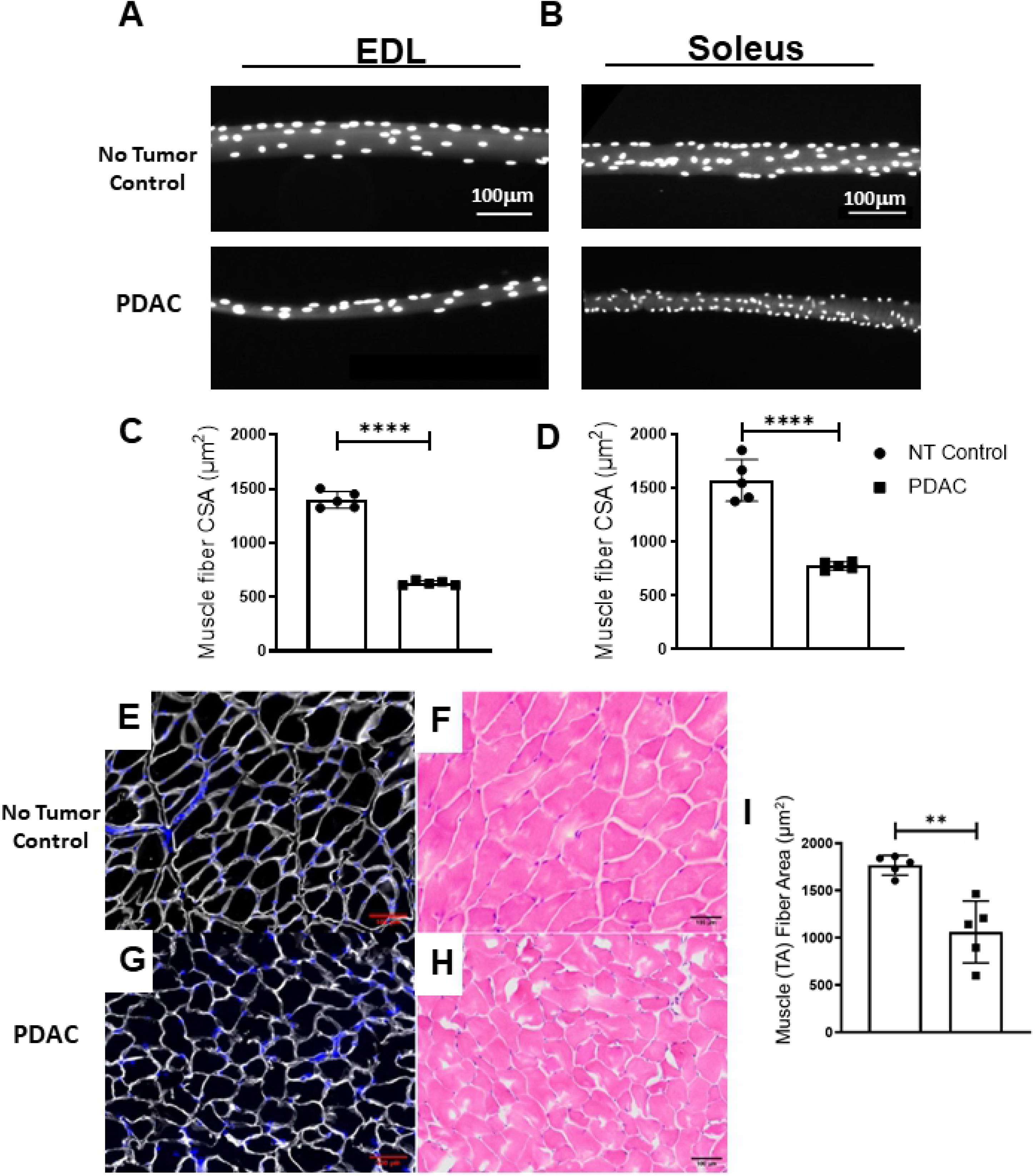
Histologic confirmation of skeletal muscle fiber atrophy in terminal mice with PDAC. Following euthanasia, the EDL and SOL muscles were harvested from PDAC baring and NTC mice and stained with DAPI for calculation of CSA (100 fibers per mouse). TA muscles were dissected and weighed. After which, TA muscles were frozen and stained with Laminin and H&E, and CSA was measured. Photomicrographs (10x) show representative DAPI stained (A) EDL and (B) SOL muscle fibers at the time of sacrifice from NT and PDAC baring mice (n=5), with quantification of (C) EDL and (D) SOL CSA (mean ± SD ****p<0.0001). Images (10x) of laminin and H&E stained TA cross-sections of (E,F) NTC and PDAC (G,H) are shown, respectively, along with quantification of TA (I) CSA (μm^2^) (n=5) and are presented with the mean ± SD (**p<0.01).

## Discussion

To better understand the mechanisms of PDAC-related SMW and identification of treatable targets, there remains a need for the development of small animal models with longitudinal outcomes. Thus, we utilized in vivo DEXA scans to predict lean muscle mass in a selected region of interest (ROI) of growing and PDAC mice. The lower portion of the hind limbs were selected as the ROI to omit the mass of the tumor from analysis. Using this method of analysis, we found strong, significant correlations between DEXA predicted lean mass and whole body and TA weight, in both the growing and PDAC mice. These findings suggest that DEXA can be used as a longitudinal outcome measure of PDAC-related SMW in a murine model.

Because of its high reproducibility and accuracy for measurement of lean body mass, DEXA has emerged as one of the most promising longitudinal assessment tools for the direct measurement of lean mass and diagnosis of sarcopenia (18). Currently, DEXA is widely used in clinical studies to diagnose sarcopenia, and is recognized as the gold standard (18).

Using a DEXA instrument for small animals to longitudinally quantify skeletal muscle mass in a cohort of PDAC mice (n=5), we found a significant reduction in skeletal muscle mass compared to their NTC littermates. This decrease in skeletal muscle mass was concomitant with “failure to thrive,” and subsequent mandatory sacrifice of the animals. TBW was not different between groups at any of the time-points. Therefore, identification of the onset of SMW may be vital for the survival of animal models and patients with cancer, if it can be halted at that time. Clinical researchers agree that once muscle loss reaches a point of clinically obvious detriment, it is irreversible (15).

Indeed, the SMW and cachexia that are experienced by patients with cancer have been defined on a continuum that begins with pre-cachexia and ends with refractory cachexia, which unfortunately cannot be relieved (15, 24). Therefore, it is important to understand when these disorders begin during disease progression and the mechanisms involved when considering viable treatment options. To our knowledge, this is the first study to assess the validity of the syngeneic KCKO-luc cell orthotopic model of PDAC as a model of PDAC-associated SMW, and the utility of longitudinal DEXA analysis to assess SMW in tumor bearing mice. The longitudinal quantification of lean mass proved to be predictive of “failure to thrive,” which was an indication for euthanasia. Thus, the diagnosis and attenuation of PDAC-associated SMW in this model can ultimately translate to improving the quality of life and survival in patients with cancer.

In addition, patients who experience SMW have a lower tolerance for treatment and higher drug toxicity (18, 25, 26), which often results in treatment discontinuation. Treatment-associated toxicity in these patients may be the result of improper drug dosing regimens that are based on antiquated methodology (18). Recent clinical evidence suggests the use of body composition measurement to improve drug dosing, reduce toxicity, and establish early intervention in patients with cancer (7, 9). Therefore, we conclude that the use of DEXA as a tool to characterize SMW to improve dosing is an area that warrants further study. Recent research reports that SMW is independently prognostic of lower survival in patients with gastrointestinal cancers (6). Martin et al.(27) showed similar results in a large cohort of patients with cancer. This study concluded that survival in sarcopenic patients was 1/3 of their non-sarcopenic counterparts, regardless of body weight. In addition, increases in lean mass during neoadjuvant treatment (NAT) has been shown to be independently associated with progression to resection surgery in patients with PDAC (9), even when these patients experienced loss of adipose tissue. Notably, patients who experienced decreases in lean mass during or after NAT underwent surgical exploration, but not resection.

Not surprisingly, we found a significant difference in TBW of animals (with tumor), compared to the animals’ normalized weight without tumor. This finding suggests that TBW is a poor indicator of survival and general health due to the continuous growth of the tumor, which mask the loss of viable lean and fat mass. These findings are significant because they specify that the longitudinal quantification of lean mass is a more appropriate prognostic indicator than the measure of TBW in this model of murine PDAC.

## Conclusion

We have demonstrated the efficacy of DEXA as a longitudinal outcome measure of PDAC-related SMW. PDAC is a disease with an extremely high mortality rate. Survival is further decreased by the onset of SMW. Early detection and treatment of SMW in affected patients may improve quality of life and survival. Therefore, more research is needed to understand the mechanism(s) that lead to a dysregulation in myocellular homeostasis. Research of this nature requires a pre-clinical model that provides longitudinal outcome measures of body composition and disease progression. In this study, we used DEXA as a longitudinal outcome measure to assess SMW in growing mice and in an established murine PDAC model. Utilizing this technique, we were able to detect the onset of SMW which was commensurate with “failure to thrive” and was confirmed through analysis of TA and body weight and histomorphometry.

